# A sparse code for natural sound context in auditory cortex

**DOI:** 10.1101/2023.06.14.544866

**Authors:** Mateo López Espejo, Stephen V. David

**Author notes:** Corresponding author: SVD.

## Abstract

Accurate sound perception can require integrating information over hundreds of milliseconds or even seconds. Spectro-temporal models of sound coding by single neurons in auditory cortex indicate that the majority of sound-evoked activity can be attributed to stimuli with a few tens of milliseconds. It remains uncertain how the auditory system integrates information about sensory context on a longer timescale. Here we characterized long-lasting contextual effects in auditory cortex (AC) using a diverse set of natural sound stimuli. We measured context effects as the difference in a neuron’s response to a single probe sound following two different context sounds. Many AC neurons showed context effects lasting longer than the temporal window of a traditional spectro-temporal receptive field. The duration and magnitude of context effects varied substantially across neurons and stimuli. This diversity of context effects formed a sparse code across the neural population that encoded a wider range of contexts than any constituent neuron. Encoding model analysis indicates that context effects can be explained by activity in the local neural population, suggesting that recurrent local circuits support a long-lasting representation of sensory context in auditory cortex.

## Introduction

The meaning of natural sounds and other sensory stimuli depends on the context in which they occur. Natural sounds are characterized by diverse temporal dynamics, like amplitude modulation at different rates. In behaviorally relevant sounds like speech, these dynamics span a range of timescales, from tens of milliseconds for phonemes, to hundreds of milliseconds for syllables and longer times for words, phrases and so on (Chomsky & Halle, 1968). Keeping track of temporal information over these diverse timescales is critical for computation and discrimination of important sound features (Norman-Haignere et al., 2022), such that altering the temporal content of speech alone, can impair its comprehension (Albouy et al., 2020).

Neurons in the auditory cortex respond selectively to spectro-temporal features of ongoing sound. Prior studies that characterized neural sound encoding with the spectro-temporal receptive field (STRF) and related linear-nonlinear models indicate that neurons in auditory cortex typically integrate information over 50-100 ms (deCharms et al., 1998; Klein et al., 2000). However, model-free analysis has shown that neurons in the auditory cortex often integrate sound information over longer periods. In primary auditory cortex, both subthreshold potentials (Asari & Zador, 2009) and spiking activity (Asokan et al., 2021) can be modulated by stimuli more than 1 second in the past. The average integration time also increases in non-primary auditory areas (Atiani et al., 2014; Norman-Haignere et al., 2022), suggesting that contextual effects on auditory coding increase hierarchically as information travels through auditory cortex.

Several neuronal and synaptic mechanisms that can contribute to temporal integration have been elucidated (Silver, 2010). Feedback signals from local inhibitory neurons have been implicated in producing enhanced response to oddball versus repeated sounds, known as stimulus-specific adaptation or SSA (Natan et al., 2015; Ulanovsky et al., 2003; Yarden et al., 2022). Stimuli can also differentially induce short-term synaptic plasticity, which can last many 10s to 100s of milliseconds, potentially modulating responses to subsequent stimuli on that timescale (David & Shamma, 2013; Lopez Espejo et al., 2019).

This neuronal integration is likely to be amplified and modulated by circuit and population dynamics (Buonomano & Maass, 2009), thus extending the computational and representational capabilities of a whole population beyond that of its constitutive neurons. Recent developments in silicon multi electrode arrays (Du et al., 2011; Steinmetz et al., 2021) have shed light on sensory representations at the population level. Among other phenomena, this approach demonstrates the existence of sparse codes in touch (Lyall et al., 2021), the dimensionality of representations in vision (Stringer, Pachitariu, Steinmetz, Carandini, et al., 2019), and a distributed encoding of time (Runyan et al., 2017). The characteristics of temporal integration in population of neurons across different auditory cortex regions, and the strategies taken by these populations to represent natural stimuli remain open questions.

While activity in auditory cortex can show long-lasting modulation by sensory context, most previous work has focused on a small number of stimulus conditions, and the specificity of these effects across the cortical population is unclear. To gain a better insight into the mechanisms underlying temporal integration, we used linear microelectrode arrays to record the activity of multiple neurons in primary (A1) and secondary (dPEG) fields of ferret auditory cortex and quantified the influence of recent stimuli on the response to ongoing natural sounds. We observed effects of sensory context lasting up to several hundred milliseconds in both areas, with a tendency toward stronger and longer-lasting effects in dPEG. Individual neurons tended to be sensitive to a small number of contexts, but the aggregate population activity formed a sparse representation tiling a much larger space of sensory context. Using encoding model analysis, we determined that local population dynamics can account for these long-lasting context effects, which cannot be explained by a traditional STRF model.

## Results

### Responses of neurons in auditory cortex to natural sounds are modulated by sensory context

To measure the effects of sensory context on neural coding of sound, we recorded single-unit neural activity in auditory cortex (AC) of awake, passively listening ferrets during the presentation of sequences of 1-s natural sound samples. Activity was recorded from neurons in primary auditory cortex (A1) and a secondary auditory cortical field (dorsal peri-ectosylvian gyrus, dPEG). Sounds were presented repeatedly and in varying order, so that the neural response to the same probe sound was recorded following many different contexts, defined as the immediately preceding sound (Figure 1A). Neural activity was recorded from linear microelectrode arrays so that the activity of tens of single units were collected simultaneously (Figure 1B, 1724 units, 64 recording sites, 5 animals).

**Figure 1.**
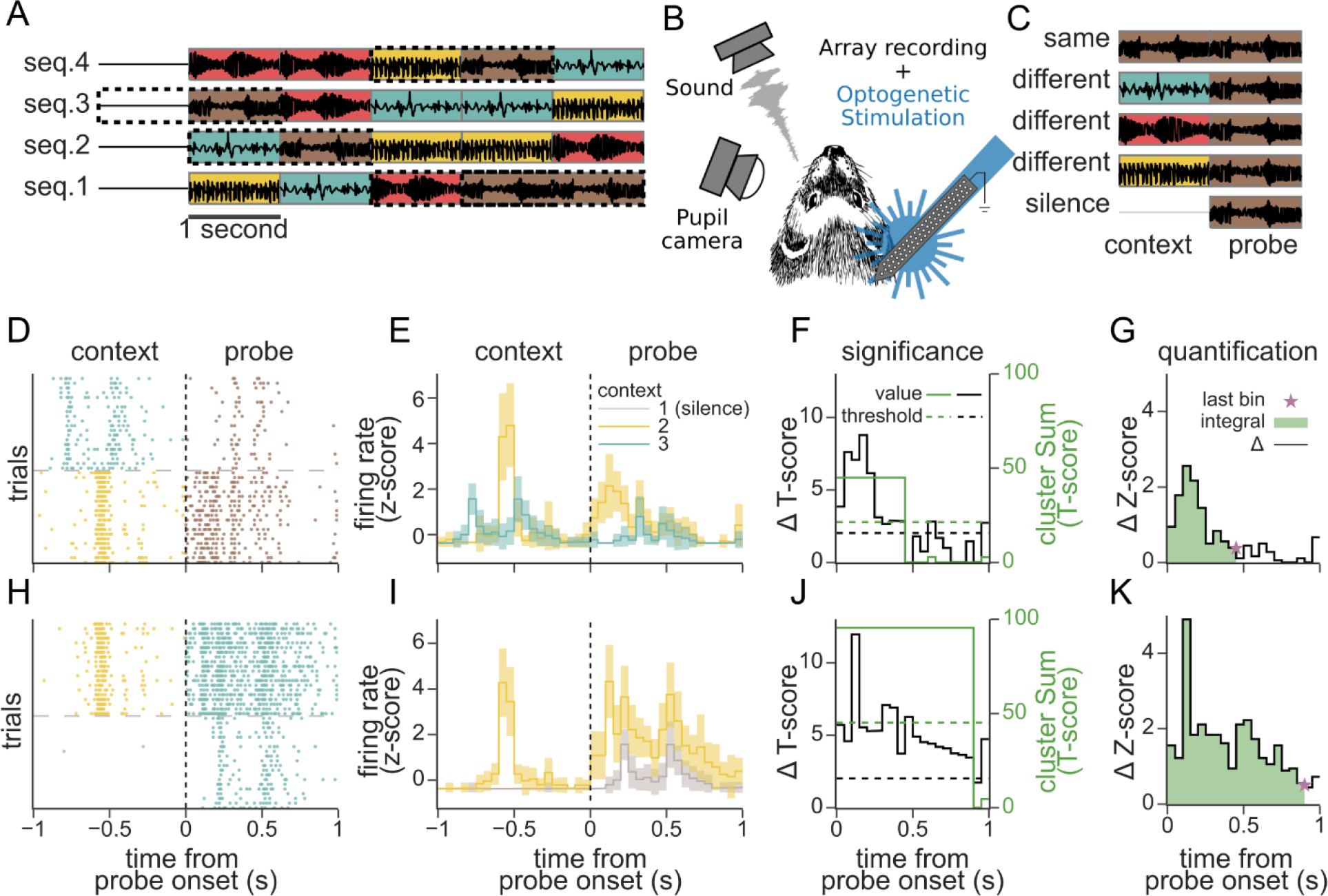
Effect of preceding sensory context on the response to a probe stimulus. **A.** Example stimulus sequences composed from four 1-second natural sounds. Sounds were ordered such that in a full stimulus set every different sound (indicated by color) followed every other sound, silence, and itself exactly once. **B.** Sounds were presented to passive listening ferrets while recording from the auditory cortex with a multi electrode array. Pupillometry was used to monitor arousal state, and optotagging of GABAergic inhibitory interneurons was performed in a subset of experiments. **C.** To assess context effects, we compared the response to one probe sound (brown, dotted boxes on a.) following each context: different sounds, silence, or the same sound. **D.** Raster of an example single unit response to 20 repetitions of the same probe (brown) after two different contexts (teal and yellow). **E.** Average peri-stimulus time histogram (PSTH) response for the raster in C (shading: 1 SEM). Line color indicates context identity. **F.** Quantification of context effects. A T-score between the probe response after two contexts was calculated for every time bin (solid black line). Clustered T-scores over a threshold (dashed black line, α=0.05) were summed (solid green line). Significance of cluster scores was determined through a shuffle test (dashed green line, α=0.05). **G.** The magnitude of significant context effects (Δ firing rate, black solid line) was quantified as its amplitude (green area under the curve, 0.559 Z-score*s) and duration (last significant bin, purple star, 450ms). H-K. Raster and PSTH response of the same neuron for a different instance of contexts and probe, plotted as in D-G (amplitude: 1.508 Z- score*s, duration: 900ms).

To measure the contextual integration window of a neuron, we computed the difference in response to a probe sound following two different context sounds (Figure 1D, H). The time-varying difference between probe PSTHs following two different contexts defined a context effect profile. We computed a T-score for each time bin (Figure 1F, J). Significant differences across multiple consecutive timepoints were identified using a cluster mass method (Maris & Oostenveld, 2007). For each neuron, we performed this comparison for each contextual instance, that is, each distinct pair of contexts preceding each probe sound. Data collected with 4 distinct sounds produced 40 context-probe instances (including silence as a context but not a probe), and data with 10 distinct sounds produced 550 context-probe instances.

We recorded the activity of 1724 AC neurons, yielding a total of 502,537 instances of context pair, probe, and neuron. In total, only 9.14% (n=45,905) of all these contextual instances showed significant effects. However, 71.5% of all neurons showed significant effects for at least one contextual instance (n=1233/1724, p<0.05, multiple comparisons correction). Because effects were highly variable within and across neurons, we analyzed each contextual instance as a distinct data point.

### Amplitude and duration of context effects varies across neurons and cortical field

Context effect profiles tended to be strongest immediately following probe onset and then decay over time (Figure 1E, H). These dynamics are consistent with the neuron having a finite integration window (Asari & Zador, 2009; Atiani et al., 2014; Norman-Haignere et al., 2022). However, the time-course showed great diversity across contextual instances and was often non-monotonic in its decay. For example, some modulation profiles had multiple peaks and valleys (Figure 1F, J). We therefore used a non-parametric approach to quantify their amplitude and duration. Amplitude was defined as the integral of the absolute delta firing rate (∫ |Δ Z-score|), across the probe response time bins with a significant difference between contexts, as identified with the T-score cluster mass test. Duration was defined as the last significant bin (Figure 1G, K).

The amplitude and duration of context effects were distributed unimodally and were correlated with each other across the neural population (Figure 2A. *r*=0.479, *p*=0, Pearson’s correlation). Some context effects lasted only briefly after probe onset and therefore had relatively small amplitude, while long lasting effects generally had greater overall amplitude. However, there were many examples of long lasting, low amplitude effects due to late onset of the contextual effect. Across all significant contextual instances in AC, amplitude and duration were both highly variable (amplitude: 0.236±0.189 Z-score*s; duration: 249.25±208.70 ms, mean ± std). In many instances (n=667, 1.4% of all significant instances), the context effects spanned the entire probe duration (1s), suggesting that context effects can last more than 1 second, consistent with previous reports of AC integration windows in other preparations (Asokan et al., 2021; Norman-Haignere et al., 2022).

**Figure 2.**
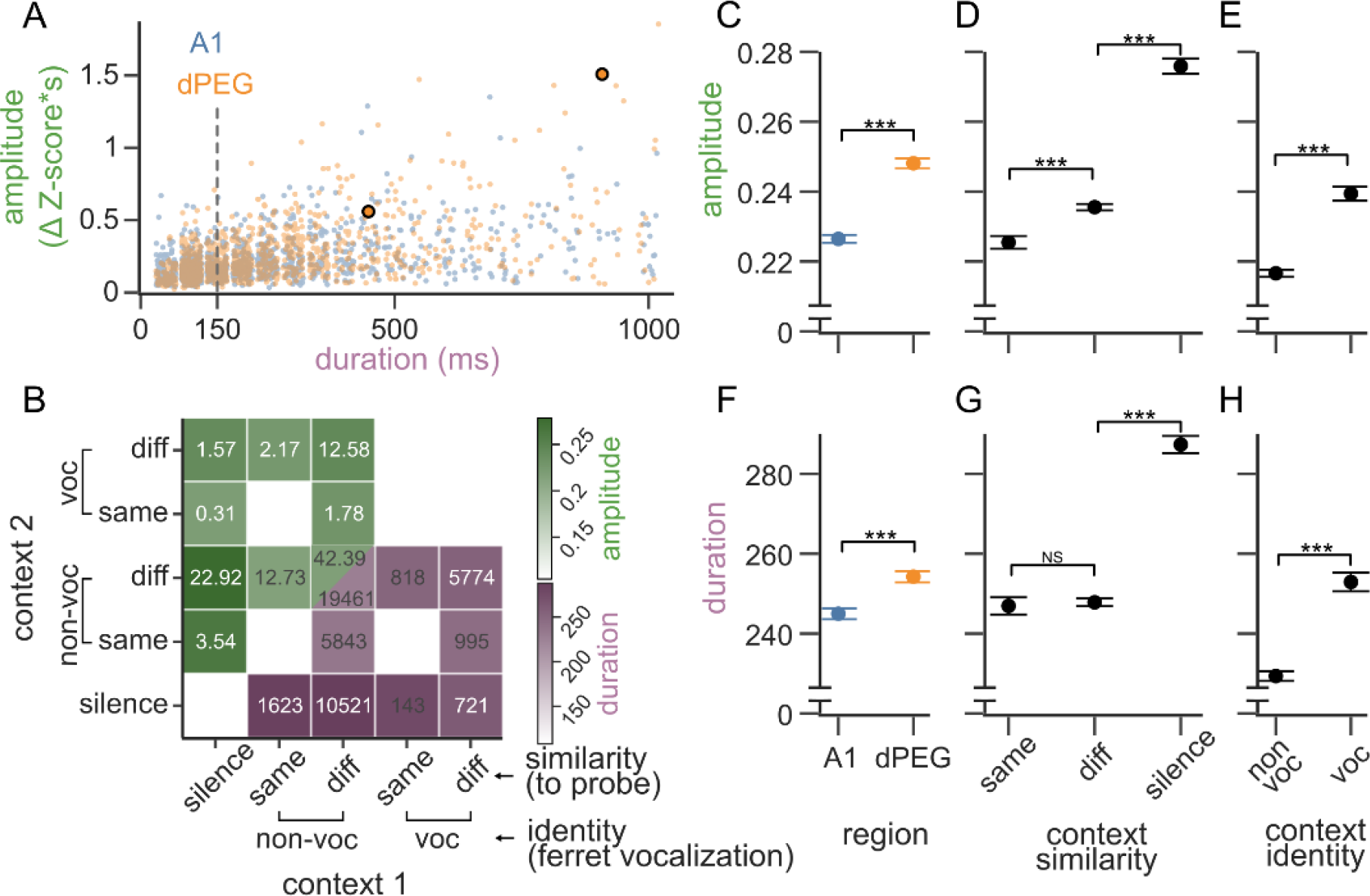
Context effect magnitude across cortical regions and context types. **A.** Scatter plot of context effect amplitude and duration for each contextual instance, *i.e.*, each combination of context pair, probe, and neuron. The two examples from Figure 1 are highlighted. Dot color indicates whether neurons were in primary (A1, blue) or secondary auditory cortex (dorsal peri ectosylvian gyrus, dPEG, orange). For clarity, duration values have been jittered from their discrete 50 ms bins, and the data was subsampled to 1000 instances from each brain region (A1: n=24711 instances, n=709 neurons. dPEG: n=21195 instances, n=523 neurons). Dashed gray line indicates the maximum temporal integration window typically observed for LN STRFs in A1. **B.** Matrix comparing mean amplitude (green, upper triangle) and duration (purple, lower triangle) of context effects for each pair of context groups (contexts 1 and 2, x and y axes). Contexts were grouped according to their identity (non-vocalization, ferret vocalization) and similarity to the probe (silence, same or different). Due to stimulus design and sound selection, some context pairs were more or less frequent (proportion and count indicated in upper and lower triangles). Others were never presented and are blank in the matrix (*e.g.*, silence-silence). **C.** Mean context effect amplitude as a function of brain region (mean and SEM, Kruskal Wallis p<0.001). **D.** Mean contribution of a context to effect amplitude based on its similarity to the probe: same, different, or silence. **E.** Mean contribution to effect amplitude of contexts identity, i.e., ferret vocalization versus non-vocalization. **F-H.** Breakdown of mean context effect duration by brain area and context category, plotted as in C-E. For comparisons in C-H: Kruskal Wallis nonparametric ANOVA with Dunn post hoc test with Bonferroni correction, p<0.001 for all comparisons except Dunn post hoc test for same-vs-diff duration, p=0.56.

Compared to A1, neurons in dPEG have complex receptive fields, associated with longer response latencies and longer temporal integration windows (Atiani et al., 2014; Bizley et al., 2005; Norman-Haignere et al., 2022). Consistent with these previous observations, dPEG neurons showed longer-lasting context effects than A1 on average (duration, A1: 244.97±1.36, dPEG: 254.25±1.40 ms, mean ± SEM; p<0.001 Kruskal Wallis test. Figure 2C). In addition, the amplitude of context effects was consistently larger in dPEG (A1: 0.23±0.001, dPEG: 0.25±0.001 Z-score*s, mean ± SEM; p<0.001 Kruskal Wallis test. Figure 2F). This result suggests that the relative weight of past memory versus current sensory response differs between areas, in addition to the duration of the integration window.

### Magnitude of context effects depends on context category and ethological relevance

We speculated that variability in the magnitude of context effects could result from a mechanism related to stimulus-specific adaptation (Carbajal & Malmierca, 2018; Ulanovsky et al., 2003). When the preceding context is silence, a probe would be a novel stimulus to which responses are not adapted and are therefore salient (onset response). In contrast, a probe that is a repeat of the context sound would be one for which responses are already adapted and thus weaker, similar to what is observed in stimulus-specific adaptation. We also speculated that the salience of context effects might be tied not only to novelty of a stimulus, but also to its ethological value. Thus, we hypothesized that species-specific ferret vocalizations might produce different context effects than the other natural sounds.

To evaluate these dependencies, we grouped contexts according to two factors, their identity and their similarity to the subsequent probe, yielding five categories: *silence*, *non-vocalization/same* as probe, *non-vocalization/different* from probe, *vocalization/same* as probe, and *vocalization/different* from probe (Figure 2B). We then used multivariate regression to quantify the relative contribution of each category to the probe response. In this model, the context effect for a single instance was a weighted sum of the identity/similarity factors of each context making up the instance, and the interaction between the factors. We quantified the effect of each context category as the percent change in the probe response relative to a baseline, defined as the most common context group, *non-vocalization/different*. As expected, amplitude and duration were largest when *silence* was one of the contexts (amplitude: +17.45%, p<0.001; duration: +16.39%, p<0.001; change relative to *non-vocalization/different*, T-test). Effects were also increased when one of the contexts was a vocalization (*vocalization/different* versus *non-vocalization/different*, amplitude: +2.08%, p=0.041, duration: +2.08%, p<0.006, T-test). Conversely, amplitude (but not duration) was lower when one context was the same sound as the probe (*non-vocalization/same* versus *non-vocalization/different*, amplitude: -4.04%, p<0.001; duration: -0.11% p=0.918, T-test). The regression model also revealed a strong interaction between *silence* and *vocalization/different* context groups. Both of these contexts tended to have a strong effect on the probe, and thus the difference between them was relatively small (interaction: amplitude: -18.44%, p<0.001, duration: -12.20%, p<0.001, T-test). No other interaction was significant.

The dependencies described above incorporate multiple aspects of the context-probe triad. In addition, we performed separate comparisons on the effects of context similarity (*silence*, *same* or *different* from probe) and of context identity (*vocalization* or *non-vocalization*) to elucidate their independent contributions. Instances with both *silence* and *vocalization* as contexts were excluded to eliminate their confounding interaction. We confirmed the increase in amplitude and duration of effects for instances including *silence* contexts (Figure 2D, G, *different* vs *silence*: amplitude: 0.24±0.0009 vs 0.28±0.0022 Z-score*s, p<0.001; duration: 247.83±0.98 vs 287.33±2.16 ms, p<0.001; mean ± SEM, Dunn post hoc), and the decrease in effect amplitude (but not duration) when one context was the *same* as the probe (*same* vs *different*: amplitude: 0.23±0.0018 vs 0.24±0.0009 Z-score*s, p<0.001; duration: 246.92±2.16 vs 247.83±0.98 ms, p=0.563; mean ± SEM, Dunn post hoc). This comparison also confirmed the increases in effect amplitude and duration in *vocalization* contexts (Figure 2E, H, *non-vocalization* vs *vocalization*: amplitude: 0.22±0.0010 vs 0.24±0.0020 Z-score*s, p<0.001; duration: 229.37±1.20 vs 252.92±2.33 ms, p<0.001; mean ± SEM, Kruskal Wallis test).

Probe identity may also independently impact context effects. When *vocalizations* were used as probes, they reduced the amplitude (but not the duration) of context effects (Supplementary Figure 1). This observation suggests that probe responses to behaviorally salient vocalizations may be resistant to contextual modulation, permitting a more faithful representation of stimuli with high behavioral relevance.

Arousal state, measured by pupil size, has been shown to impact sound-evoked activity in cortex (McGinley et al., 2015; Schwartz et al., 2020). Salient stimuli can increase arousal, and we considered the possibility that some context effects might emerge indirectly from differential changes in arousal following more or less salient stimuli. Pupil size was positively correlated with an overall increase in evoked firing rate (Supplementary Figure 2). Early context effect size was also positively correlated with pupil size, but it was negatively correlated at later timepoints. The decrease may serve to increase fidelity of representation of the current probe relative to the preceding context in high arousal states. Overall, pupil-related changes were small, indicating a relatively weak contribution of arousal to context effects.

### Auditory context is represented in a sparse, distributed code

We presented each neuron with multiple combinations of contexts and probes, which together comprised a context space (40 and 550 combinations for datasets using 4 and 10 natural sound samples, respectively). Not all sound combinations elicited significant context effects; therefore, individual neurons represented a limited extent of the context space (Figure 3 A, 4-sound examples). On average, a neuron covered only 11.2 ± 0.374% (Figure 3C, Mean ± SEM) of the space, though neurons in dPEG responded to more contexts than those in A1 (Figure 3C, *single-neuron*: A1: 9.17±0.43, dPEG=14.04±0.64, p<0.001; mean ± SEM, Mann Whitney U test). However, different neurons from the same recording showed effects in distinct regions of the context space, generating a sparse code for context effects.

**Figure 3.**
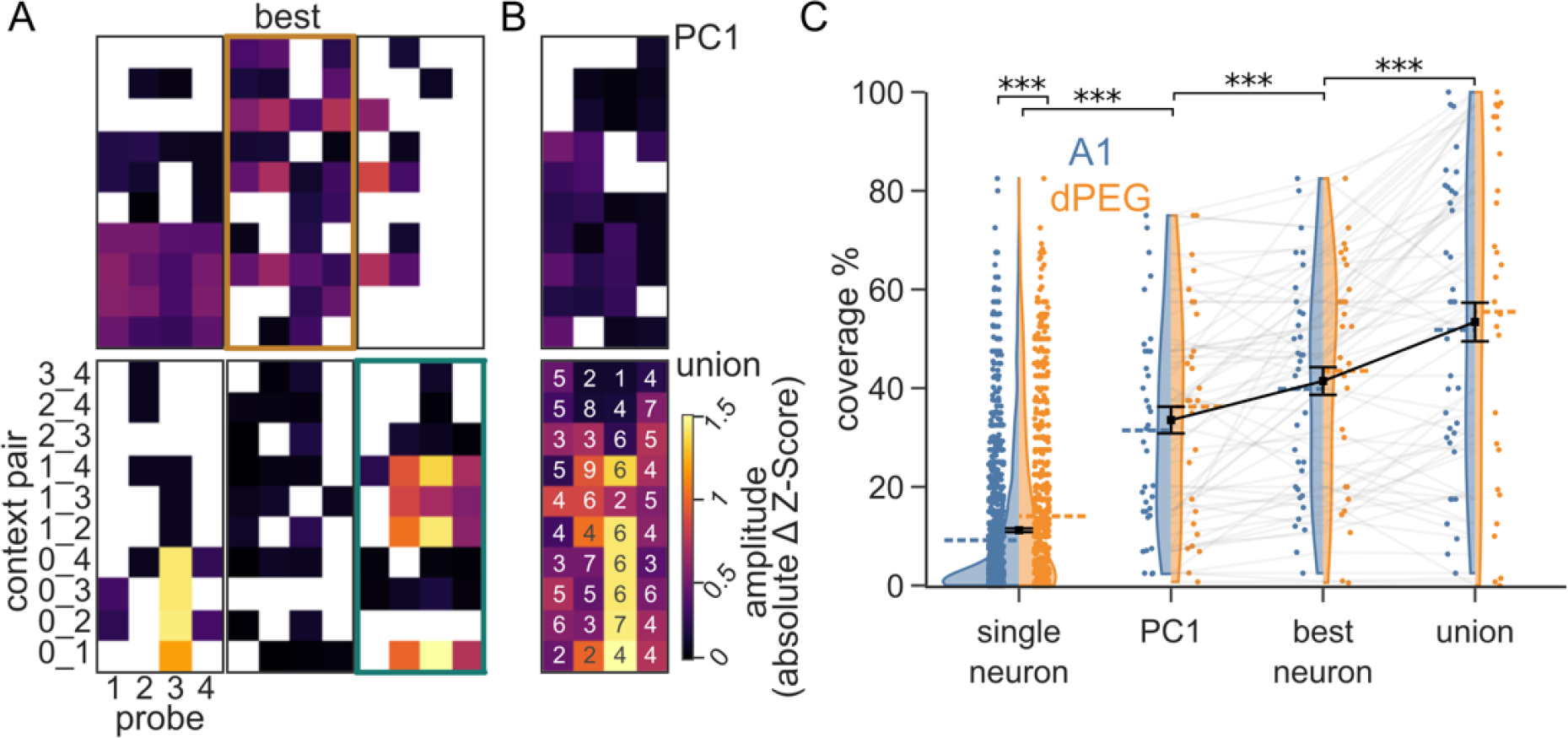
Context is represented sparsely in AC populations. **A.** Heatmaps show contextual coverage, *i.e.*, amplitude of context effects across all context instances (4 distinct probes/10 context pairs) for 6 simultaneously recorded neurons. Y-axis indicates indices of context pair sounds, with 0 denoting silence. Only significant effects are colored. The number of significant instances varies from 9 to 29 across the examples shown. The neuron with the greatest coverage (29/40 significant instances) is denoted best. Neurons outlined in orange and teal are used as examples in Figure 4. **B.** Illustration of two methods for measuring context effects for activity pooled across a recording site. Context effect amplitude was measured for the first principal component of population activity across the entire recording site (top, n=33). The union of context effects was computed across all neurons in the recording site (bottom, numbers denote neurons with significant effects per instance after Bonferroni correction for number of neurons). **C.** Distribution of percent contextual coverage for single neurons (A1 n=1006, dPEG n=716 neurons), and pooled activity for each recording site (A1 n=36, dPEG n=28 sites. Blue: A1, Orange: dPEG. Dashed colored lines indicate the region’s average. Black squares and error bars show mean and SEM across both regions). For comparison between brain regions: Wilcoxon signed-rank test, significant only for single neurons, p<0.001. For comparisons against single neurons and between site groups: Mann Whitney U and Wilcoxon signed rank tests respectively, both with Bonferroni corrections, all comparisons significant, p<0.001.

Sparse codes are reported widely across sensory systems (Lyall et al., 2021; Olshausen & Field, 1996; Zhang et al., 2019) and hypothesized to provide multiple advantages in associative learning, storage capacity, energy efficiency and facility to read out the encoded information (Beyeler et al., 2019; Olshausen & Field, 2004). Because we recorded the activity of multiple neurons simultaneously, we could measure context information present in the population activity across an entire recording site. We used two complementary methods. First, we measured context effects for the first principal component of the population activity (*PC1*). Second, we computed the *union* of context effects across all the neurons in the site (Figure 3B). The *PC1* and *union* approximate the information available to optimal linear- and nonlinear decoders, respectively, downstream from auditory cortex. In both cases, context coverage was well above the average for *single neurons*, consistent with a sparse, distributed code (Figure 3C, p<0.001 for all comparisons, Mann-Whitney U test with Bonferroni corrections). However, only the *union* showed greater coverage than the *best neuron* in each site (Figure 3C, *PC1:* 33.53±2.70%, *best neuron:* 41.44±2.79%, *union*: 53.40±3.91%, mean ± SEM, p<0.001 for all comparisons, Wilcoxon signed rank test with Bonferroni corrections).

Sparse codes support an easier readout of sensory information by downstream decoders (Beyeler et al., 2019). To quantify the information available in the population activity, a support vector machine was trained to predict the identity of probe or contexts at every time point after probe onset. Over time, decoder accuracy for the context decreased, and accuracy for the probe increased (Figure 4C). These dynamics are consistent with context information persisting over a period after context offset, simultaneous with accumulation of information about the current probe. To test if sparse, neuron-dependent code for context facilitated readout, we transformed the population activity such that context effects were equally (densely) distributed across all the neurons in the population and matched the original total magnitude of the context effect across the population. From a geometric state-space perspective, this can be visualized as shifting of the context dependent activity to the diagonal of equal neuronal activity (Figure 4D), while preserving single trial variability and the context-independent probe response. The consequence of encoding context with a dense representation was readily observed in the PSTH difference between contexts, which became identical for all neurons in a site (Figure 4E), but with smaller amplitudes relative to the original neurons that showed effects (Figure 4B). The dense transformed responses severely reduced the context decoder performance, while leaving the probe decoder performance unchanged (Figure 4F). Because decoder performance depends on the number of contexts in the dataset, we focused on the data collected with 10 sound stimuli. However, we observed a very similar pattern when we analyzed data collected with 4 sound stimuli (Supplementary Figure 3).

**Figure 4.**
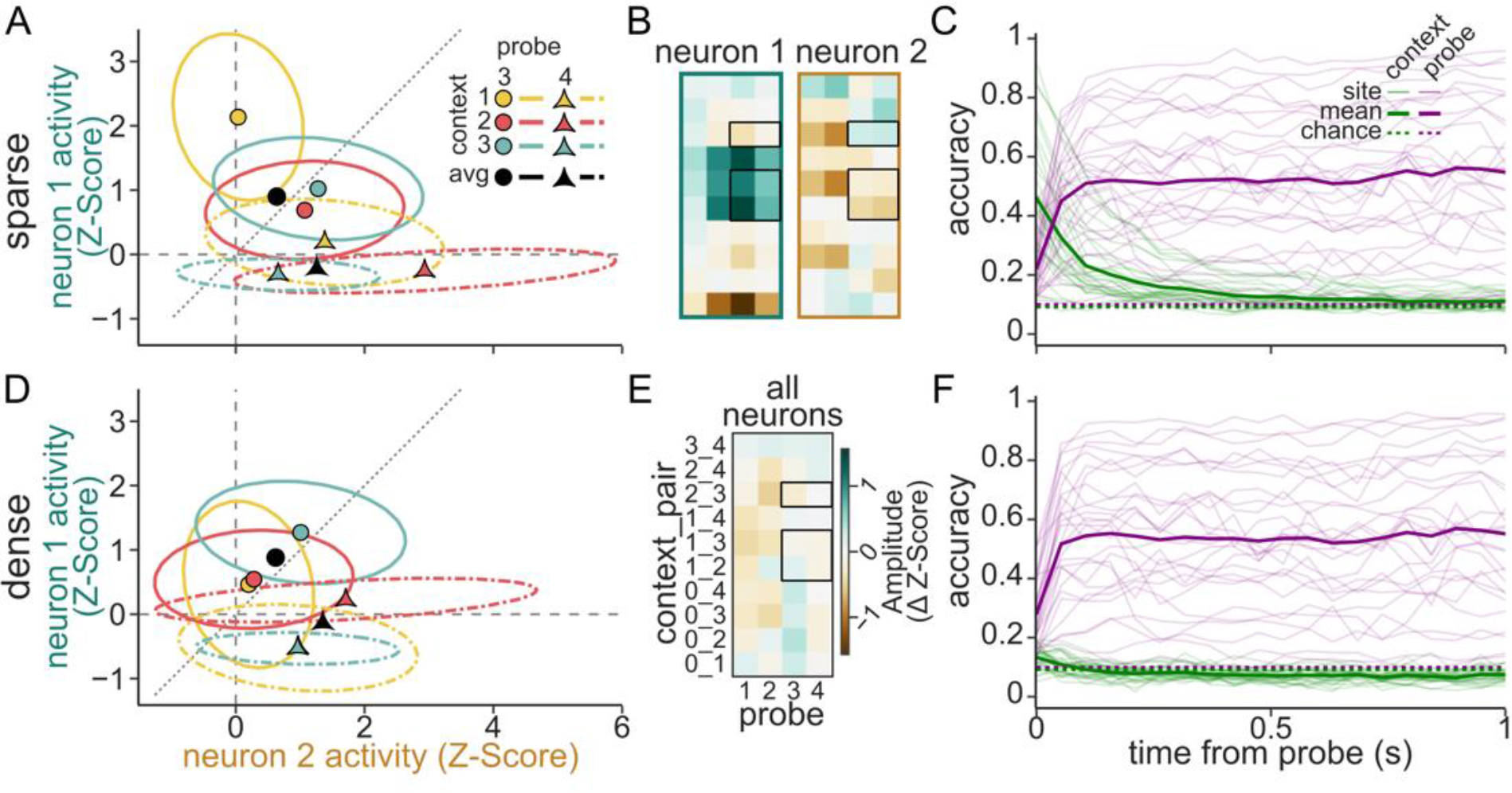
Sparse population code supports linear decoding of auditory context. **A.** Example response of two neurons (x- vs. y-axis, colored as in Figure 3A) to two probe stimuli (shape, line type) following 3 contexts (color) at a single time bin from the original data. Colored markers indicate the trial average response, and ellipses represent one standard deviation of across trials. Black markers indicate the average probe response across all contexts, *i.e.*, the context-independent response. The amount of overlap between two ellipses corresponds to the decoding accuracy between the corresponding stimulus pair. **B.** Heatmaps show PSTH response difference for each contextual instance for the two example neurons (plotted as in Figure 3A). Black boxes highlight instances in A. **C.** Support vector machine accuracy at decoding context (green) and probe (purple) identity at each timepoint during probe presentation. Thin lines show performance of individual sites (n=28 sites exposed to 10 different sounds) and thick lines indicate average performance. Chance performance is indicated with dotted lines. **D.** Simulated responses of the same neurons as in A after imposing a dense representation of context, in which context effects have the same overall magnitude but are constant for each neuron. Individual context effects differ from A, but probe responses are the same. **E-F.** Context coverage and decoding accuracy for the dense context code, plotted as in B-C.

### Context effects are weaker but more common in putative inhibitory interneurons

Previous work has implicated inhibitory interneurons as having specialized roles in temporal processing of sound (Wehr & Zador, 2003), SSA (Natan et al., 2015; Yarden et al., 2022) and in sensory integration (Studer & Barkat, 2022). Thus, we hypothesized that they also play a distinct role in the representation of sensory context. We used a viral approach to express Channelrhodopsin (ChR2) selectively in inhibitory interneurons (Supplementary Figure 4) (Dimidschstein et al., 2016), which were then identified by optotagging (Figure 5A). Because ChR2 was only transduced in a subset of recordings and may not have labeled all inhibitory neurons when expressed, we also used spike width (peak-to-trough delay, PTD) to distinguish neurons as narrow spiking putative inhibitory interneurons (n=301 neurons; PTD<0.37ms) and broad spiking pyramidal cells (n=1172 neurons, PTD>0.47ms. Figure 5B, C) (Trainito et al., 2019). The narrow spike widths of the optotagged neurons were consistent with that of putative inhibitory interneurons.

**Figure 5.**
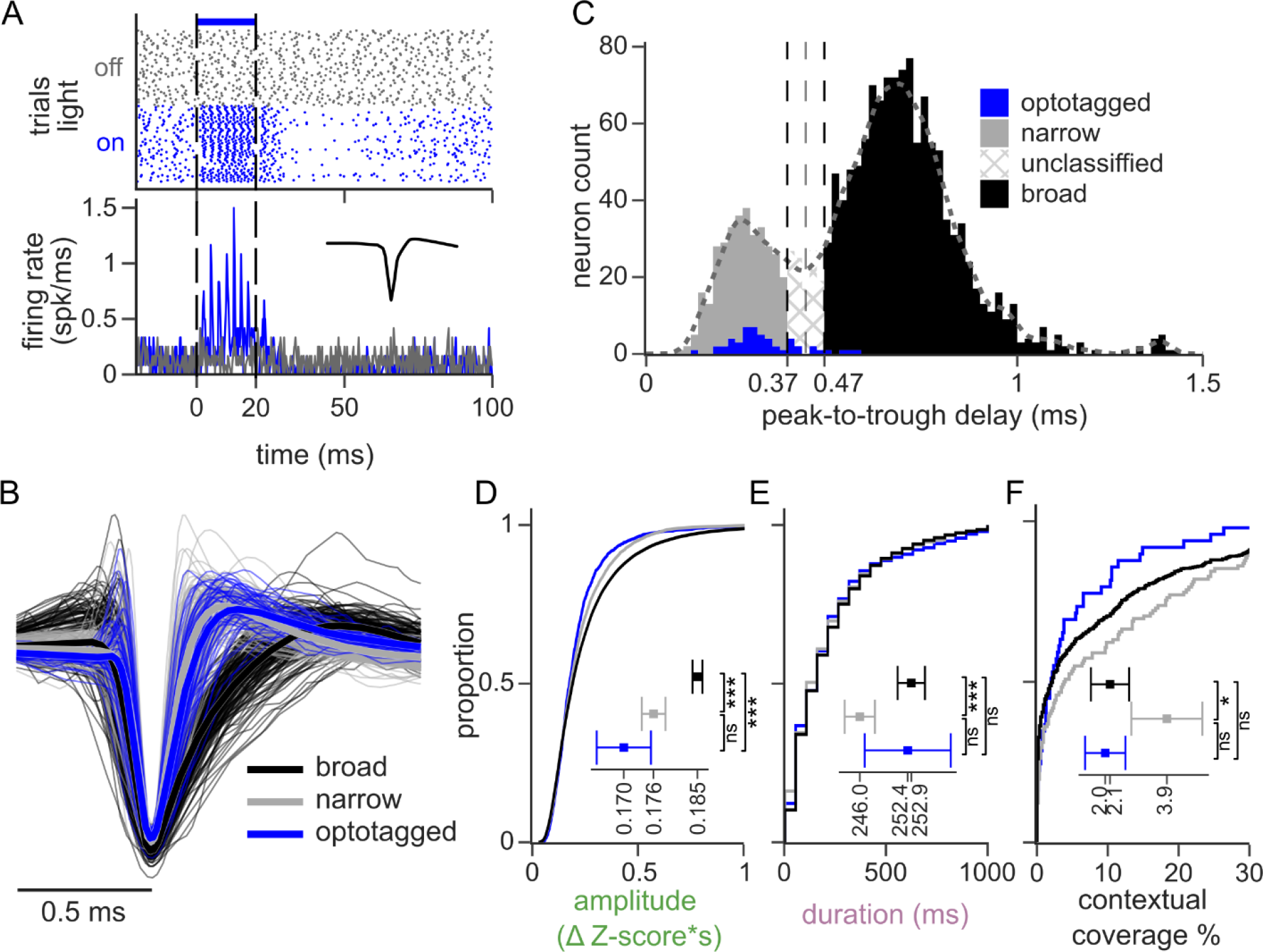
Context effects are weaker but more common in inhibitory interneurons. **A.** Example single trial responses (top) and average PSTH (bottom) of an optotagged A1 neuron to silence (gray) or a continuous 20ms light flash (blue). Vertical dashed lines show light onset and offset. The inset in the lower panel shows the average spike waveform. A neuron was classified as optotagged if it responded with sustained spiking <5ms after the light onset and showed a reliable spiking pattern across trials. **B.** Example spike waveforms (thin lines) and averages (thick lines) for neurons classified as narrow spiking (gray), broad spiking (black), and optotagged neurons (blue). Waveforms normalized to a fixed peak. For clarity, only 500 random examples are shown per color. **C.** Histogram of spike peak-to-trough delay colored by cell type: narrow-spiking (n=301 neurons, putative inhibitory, gray), broad-spiking (n=1172 neurons, putative excitatory, black) and optotagged inhibitory neurons (n=51 neurons, blue). Single units with intermediate peak-to-trough values were unclassified (0.37-0.47 ms). **D-F.** Cumulative histogram of the context effect amplitude (D), duration (E), and contextual coverage (F) for the classified cell types. Insets show the median amplitude and coverage, and mean duration, with the 100-jackknife confidence interval (ns: non-significant, *: p<0.05, ***: p<0.001. Kruskal-Wallis with post hoc Dunn test).

Narrow spiking neurons showed context effects of smaller amplitude and duration relative to broad spiking neurons (narrow vs broad: amplitude: 0.176±0.002 vs 0.185±0.001 ΔZ-score*s, median ± SE, p<0.001; duration: 246.01±2.03 vs 252.90±1.83 ms, mean ± SE, p<0.001. Dunn post hoc tests, Figure 5D, E). Despite having smaller context effects, narrow spiking neurons showed them more frequently. That is, they showed modulation over a greater proportion of context instances tested (narrow vs broad: 3.9±1.1 vs 2.1±0.5 % coverage, p=0.022, median ± SE, Dunn post hoc test, Figure **5**F). Optotagged neurons showed no difference from narrow spiking neurons (amplitude: 0.169±0.005 ΔZ-score*s, p=0.13; duration: 252.42±5.76 ms, p=1.0; coverage: 2.0±0.6 % coverage, p=0.23. mean/median ± SEM, Dunn post hoc tests vs narrow, Figure **5**D-F) supporting their classification as inhibitory interneurons. Furthermore, optotagged neurons differed from broad spiking neurons, albeit only for context effect amplitude (amplitude: p<0.001; duration: p=0.22; coverage: p=1.00, Dunn post hoc tests vs broad, Figure **5**D-F). This is likely due to the smaller effects for duration and coverage, and the reduced statistical power due to the limited number of optotagged neurons.

### Encoding model analysis indicates a role of local population activity in representing context

Finally, we used an encoding model approach to evaluate mechanisms that may underlie the observed context effects. We hypothesized several possible factors that could contribute to context effects: (i) long-latency receptive field integration times, (ii) feed-forward adaptation to sensory inputs, (iii) and modulation by local neural population activity. To evaluate the role of these different mechanisms, we fit a set of generalized linear models (David, 2018; Thorson et al., 2015) that successively incorporated terms to account for their effects (Figure **6**A). The baseline spectro-temporal receptive field (*STRF*) model described the activity of a neuron based only on the sound spectrogram, providing a standard model of sound encoding in AC (Atiani et al., 2014; deCharms et al., 1998). To account for possible adaptation effects, the *Self* model included an additional input based on the past spiking activity of the neuron being modeled. To account for local population effects, the *Pop* model incorporated past activity of the simultaneously recorded neurons. Finally, the *Full* model incorporated both the neuron’s own activity from the *Self* model and the local population activity from the *Pop* model.

**Figure 6.**
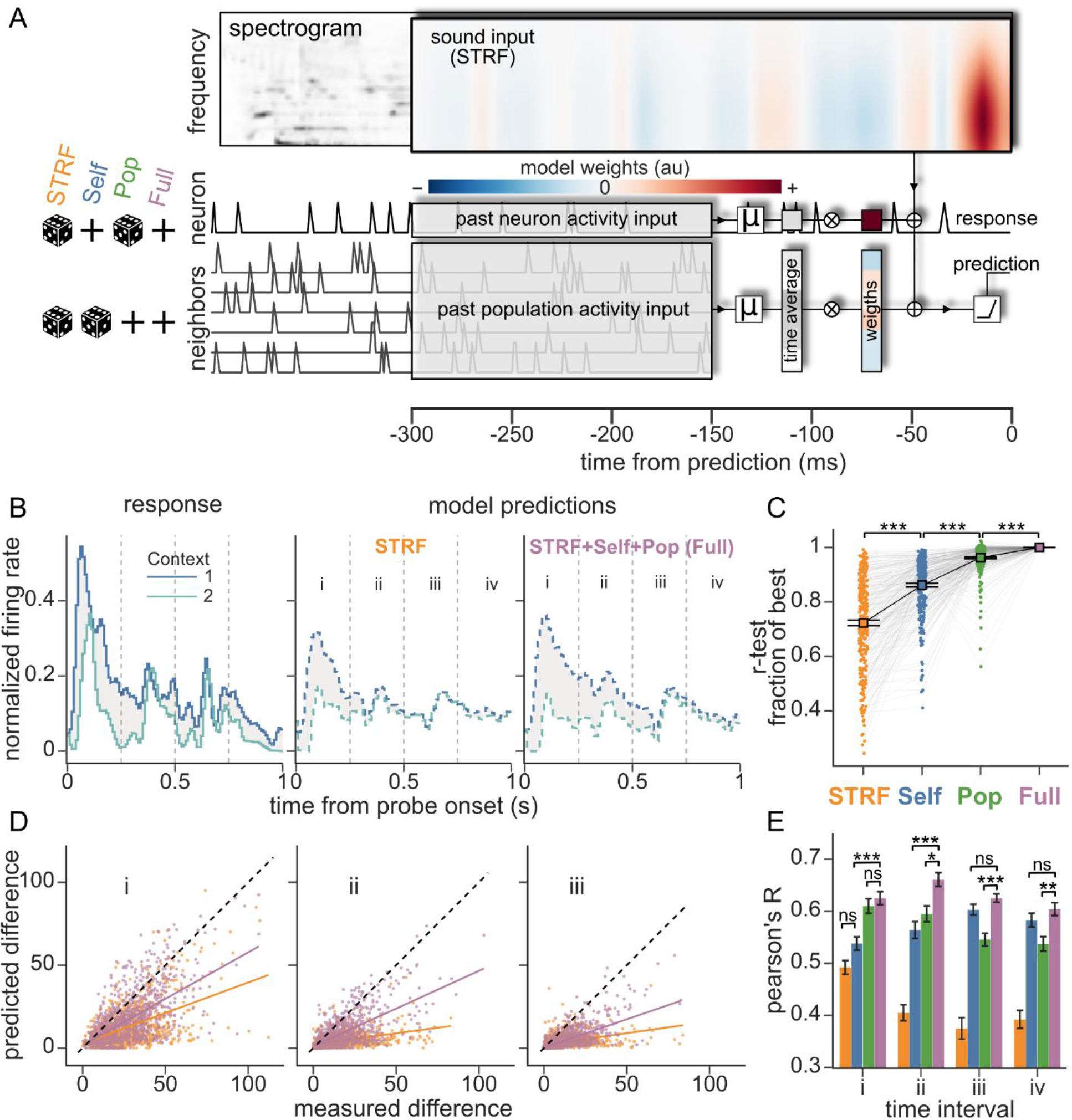
Local population activity predicts subsequent context effects. **A.** Architecture of encoding models predicting a neuron’s responses as a function of the immediately preceding sound and history of its own activity (self) and of the activity of neighboring neurons (pop). Sound information (spectrogram) was weighted with a spectrotemporal receptive field (STRF) spanning 18 spectral channels and a latency of 0-300 ms. Self and pop history were averaged over a window 150-300 ms in the past and scaled with neuron-specific weights (colored vectors). Activity <150 ms in the past was excluded to remove the confound of correlated sensory responses at shorter latency. The weighted sound and past neuronal activity were summed and passed through a rectifying linear unit (ReLU) to generate the prediction. To isolate the contribution of past neural activity to predictions, these inputs were time-shuffled (dice) or not (plus sign), defining four different models (left side glyphs. *STRF*: both shuffled, *Self*: population shuffled, *Pop*: neuron shuffled, *Full*: no shuffling). **B.** Example neuron response (left) and predictions for the *STRF* (center) and *Full* (right) models to a single probe after two contexts (blue, teal). The contextual effect (gray area) was calculated for 250ms intervals (i-iv). **C.** Quantification of prediction accuracy (cross validated Pearson’s r) for each model as a fraction of *Full* model accuracy. Colored circles connected by gray lines represent models fit to individual neurons (n=275). Black squares and error bars are the mean and SEM with a corresponding trend line (STRF vs Self: p<0.001, Self vs Pop: p<0.001, Pop vs Full p<0.001. Wilcoxon signed-rank test, Bonferroni corrected). **D.** Comparison of context-dependent difference for the actual response versus prediction by the *STRF* (orange) and *Full* (purple) models, at time intervals i (left) ii (center) and iii (right). Data wase pooled across all significant instances (n=15176) for all fitted neurons. For clarity, data were subsampled by taking 1000 random points per model. Linear regression and Pearson’s correlation coefficients were calculated over all the data (colored lines). **E.** Pearson’s correlation of predicted context effect amplitude for all models and time intervals (i-iv). Error bars are the standard deviation calculated from the 200-fold jackknifed distribution of Pearson’s r. Models were compared within each time interval. Comparisons not specifically labeled were significant with p<0.001 except A-Self vs A-Pop p<0.01, B-Self vs B-Pop p=0.8, C-Self vs C-Pop p<0.01 and D-Self vs D-Pop p=0.1 (ns: non-significance, *: p<0.05, **: p<0.01, ***: p<0.00. Student’s T-test with Bonferroni correction)

This set of models can be described as a standard STRF plus optional inputs for self- and population history. To balance parameter count between models, all models contained parameters for the addition inputs, but the inputs were temporally scrambled when the term was not included in that model. Because all models had the same number of free parameters, differences in their performance were a direct consequence of the value of the predictors, and not differences in estimation noise.

Models were fit using a separate set of natural sounds presented to the same neurons but not used to measure context effects (n=275 neurons presented with both stimulus sets). We then used the models to predict the time-varying spike rate response to the context-probe stimuli. Overall performance was calculated as the Pearson’s correlation between real and predicted activity. Model function and this performance measurement were agnostic to context and probe classification or transitions. Models incorporating past neuronal activity as predictors performed significantly better (Figure **6** C, Full>Pop>Self>STRF), indicating that the additional terms were accounting for activity that was not captured by the STRF.

To measure models’ ability to capture context effects, we compared real and predicted probe responses. Due to the deterministic nature of our models, we could not measure context effects for the predictions using the same T-Score cluster mass method as was used for the actual data. Instead, we simply measured the amplitude of the difference in predicted probe response between contexts. To discriminate the temporal profile of context effects captured by the models, we computed the amplitude of context differences separately for non-overlapping 250ms intervals in the probe response (intervals i-iv, Figure **6** B). Paralleling the overall measures of prediction accuracy, models incorporating information about history of neural activity better predicted context effect amplitude at all time intervals: The baseline STRF model captured some context effects immediately after probe onset (interval i), but it failed to capture later effects. In contrast, the full model captured later context effects, with peak performance during interval ii, between 250 and 500ms (Figure **6** D), but with significant improvement also during the later intervals ii and iv (Figure 6 E).

The Self and Pop models, which incorporated only one history term, also performed better than the baseline STRF at most time intervals (Figure 6E). However, they approached Full model performance during different time intervals. The Pop model explained most of the performance gain of the full model early (Pop vs Full, interval i, non-significant difference), but the Self model matched the Full model performance during later intervals (Self vs Full, interval iii and iv, non-significant differences). This pattern suggests distinct roles of population activity and the neuron’s own history (adaptation, plasticity), with population dynamics carrying most contextual information early on, and later being superseded by the neuron’s internal state.

## Discussion

### A sparse, distributed representation of natural auditory context

We observed effects of sensory context on cortical responses to natural sounds lasting up to several hundreds of milliseconds, consistent with previous observations (Asari & Zador, 2009; Klampfl et al., 2012). Neurons showing these effects often only did so under specific stimulus conditions, that is, only for a few combinations of contexts and probe sounds. While context effects were sparse for individual neurons, context sensitivity varied across groups of simultaneously recorded neurons. This diversity of effects supports a sparse, distributed code for sensory context at the population level. The sparse representation of context effects is reminiscent of sparse codes previously reported for sensory tuning in the visual, somatosensory and auditory cortices (Lyall et al., 2021; Olshausen & Field, 1996; Zhang et al., 2019). Sparse codes provide an efficient representation strategy, yielding independent representations that can be read out easily and robustly with a linear decoder (Beyeler et al., 2019; Rigotti et al., 2013), and our decoding analysis demonstrates the benefit of the distributed code. Context could be decoded from population activity for several hundred milliseconds after probe onset. On the other hand, simulated neurons with a dense code for context permitted much weaker decoding performance (Rigotti et al., 2013). Thus, by examining context effects across many combinations of context and probe stimuli, our observations support the idea that sparse codes are used to integrate sensory information over much longer timescales than is observed in standard models of sensory coding.

We observed a small change in sparseness between primary and non-primary cortical fields A1 and dPEG, respectively. The sparseness of context representation is decreased in dPEG. This result suggests that as more complex representations emerge along the auditory pathway, the sparse constituents of these representations in A1 are combined, becoming less sparse though mixed selectivity (Rigotti et al., 2013). However, the emerging representation may be sparse in some higher-order representational space, facilitating further processing in downstream areas.

### Magnitude of context effects reflects stimulus salience

We hypothesize that the magnitude of context effects is partially driven by the salience of both the context and probe stimuli. This hypothesis is supported by the increased effects when one of the contexts is silence or a behaviorally relevant ferret vocalization. Superficially, these are seemingly opposite cases, where the salience lies in either the onset of a probe stimulus after silence, in the former case, versus a salient vocalization context stimulus in the latter. However, we argue that both cases support an enhanced representation of the relevant stimuli through either a strong onset detection (following silence) or sustained activity (following a vocalization) that outlasts the stimulus and enables processing. The enhanced representation of salient sounds is further supported by the faithful representation of vocalizations when they occur as probes, for which context effects are smaller and therefore provide less interference to the probe response. This preferential representation of behaviorally relevant stimuli has been reported previously, though the mechanism of receptive-field expansion following learning (Elgueda et al., 2019) and immediately after engaging in a behavior (Fritz et al., 2003).

The increased response and context effects related to salient contexts (Vocalizations) or probes (sound after silence) are reminiscent of stimulus-specific adaptation (SSA) (Natan et al., 2015; Yarden et al., 2022) and show how adaptation phenomena may influence sound coding of wide-ranging natural stimuli. It follows that the mechanisms underlying SSA, including short term synaptic plasticity, distinct synaptic input, and top-down feedback (Malmierca et al., 2015), are likely to participate in generating context effects. This hypothesis is supported by the improvement of encoding models by using the neuron recent activity, a proxy for adaptation, as a predictor. Furthermore, we speculate that arousal contributes to the magnitude of context effects associated with salient stimuli, through a mechanism of response amplitude modulation (Schwartz et al., 2020). We observed high arousal significantly increased context effects early after probe onset, and decreased them later, thus enhancing the representation of salient stimuli like onsets. However, these effects had a small amplitude, and we argue that increased arousal only plays a limited role in increasing context effects and enhancing the representation of novel stimuli.

### Mechanism supporting context effects

Standard linear STRF models can describe the tuning of early auditory neurons (Aertsen & Johannesma, 1981; Escabí & Read, 2003; Kowalski et al., 1996), but they fail to capture sound-evoked activity at latencies >150 ms (Atiani et al., 2014). Thus, a more comprehensive model is required to explain the long-lasting context effects reported here. Nonlinear mechanisms related to coding of modulation rate (Joris et al., 2004; Lu et al., 2001; Sharpee et al., 2011), adaptation to stimuli (Ulanovsky et al., 2003), and noise invariance (Rabinowitz et al., 2013), might support information integration over hundreds of milliseconds required for computation of sound features related to context and expectations.

We considered encoding models that could account for some of these longer-lasting nonlinear response properties by expanding and STRF model to include the history of activity of neighboring neurons. Models incorporating either the neuron’s own history or the history of neighboring neurons were able to better account for responses to the context-probe stimuli, and they accounted for context effects on complementary timescales. The processes captured by these local activity models may reflect a combination of feedforward connectivity acting on faster time scales (Wehr & Zador, 2003), and recurrent activity acting on longer timescales (Carandini & Heeger, 2012; Oldenburg et al., 2022). The faster feedforward computations might be related to early context effects past the sharp probe onset response, needed for representing the statistics of ongoing sound and reporting their sudden change though hidden state computation (Buonomano & Maass, 2009). Finally, late context effects are likely due to recurrent activity, locally supported by inhibitory interneurons (Saha et al., 2017), and in even later context effects as consequence of long-distance feedback loops trough the midbrain (Malmierca et al., 2015) or other cortical regions (Bizley et al., 2015).

Our results also suggest that inhibitory interneurons play a specialized role in context effect representation. They show small amplitude context effects, but over a wider range of contexts. We hypothesize that the local activity pooling characteristic of inhibitory interneurons (Carandini & Heeger, 2012), increases the complexity of their receptive fields (Moore & Wehr, 2013) explaining the wider range of context effects. At the same time, activity pooling leads to averaging, which explains the reduced amplitude context effects. Furthermore, inhibitory interneurons likely play other roles in temporal integration and context representation though SSA mechanism (Natan et al., 2015; Yarden et al., 2022), supported by the distinct facilitation and depression of the synapses of PV and SOM interneurons in thalamic feedforward motifs (Tan et al., 2008).

## Methods

### Animal preparation

Adult male ferrets (aged 6-9 months) were surgically implanted with a head post to stabilize the head and enable multiple small craniotomies for acute electrophysiological recordings. Anesthesia was induced with ketamine (35mg/Kg) and xylazine (5mg/kg) and maintained with isoflurane (0.5-2%) during the surgical procedure. The skin and muscle on top of the cranium were removed and the surface of the skull was cleaned. Ten to twelve small surgical screws were placed on the edge of the exposed skull as anchor points. The surface of the skull was chemically etched (Optibond Universal, Kerr) and a thin layer of UV-cured dental cement (Charisma Classic, Kultzer) was applied over the exposed surface. Two stainless steel head posts were aligned along the midline and embedded with additional cement. Finally, cement was used to build a rim extending out from the edges of the implant. The rim served the dual purpose of holding bandages over the implant margin wounds and creating wells to hold saline over the recording sites. Once the implant was finished, excess skin around it was removed, the wound around the implant was closed with sutures and the animal was bandaged. Antibiotics and analgesics were administered as part of the post-op recovery.

After at least 2 weeks following surgery, the animals were acclimated to a head-fixed posture, during intervals starting at 5 minutes and increased 5 to 10 minutes every day. Food and liquid rewards were given during these acclimation sessions to help the animals relax under restraint. Animals were considered ready for recording when they could be restrained for more than 3 hours without signs of distress (e.g., the animals being relaxed enough to fall asleep).

### Sound presentation

Acoustic stimuli were either synthesized or drawn from a library of pre-recorded samples and presented using custom Matlab software. Digitized signals were converted to analog (National Instruments) and amplified (Crown). They were presented to head-fixed animals in a sound attenuating chamber (Gretch-Ken or Acoustic Systems), using calibrated free-field speakers (Manger) positioned at 30-deg contralateral azimuth, 0-deg elevation, and 80 cm distant from the animal.

Auditory stimuli used for measuring sensory context effects were sequences of 1-second natural sounds, which we refer to as context-probe pairs. Sequences were constructed so that each probe sound was preceded by several different context sounds. To maximize efficiency of context-probe sampling, we generated sequences of N different sounds, such that any sound acted as the probe following a preceding context, or as the context for the following sound. Each sound was also played at the beginning of the sequence, therefore acting as a probe following a silent context. Sounds were also repeated so that a probe could provide its own context (Asari & Zador, 2009).

For N different sounds, full sampling of context-probe combinations was achieved with N sequences of N+1 sounds (Figure 1A). Finding sound sequences fulfilling these conditions poses a mathematical problem known as “exact coverage”, which we solved using the dancing links algorithm (Knuth, 2000). We created sequences from N=4 or 10 1-second sound samples. In each experiment, sounds were drawn from a set of 16 natural sounds, based on their ability to drive neuronal activity in the recording site. This 16-sound set was chosen from a large library, selected for their ability to drive activity across many neurons in previous recordings in the laboratory. It contained music, speech, ferret vocalizations and environmental noise such as gravel and cash registers.

### Neurophysiological recording

The putative location of A1 and dPEG was determined during the headpost implantation surgery based on external landmarks: the posterior and medial edges of A1 falling, respectively, 13 mm anterior to the occipital crest and 8 mm lateral to the center line, and dPEG immediately antero-lateral to A1 (Bizley et al., 2005). To functionally confirm recording locations, we opened small craniotomies <1mm diameter and performed preliminary mapping with tungsten electrodes (FH-Co. Electrodes, AM Systems Amp, MANTA software (Englitz et al., 2013). We measured the tuning of the recording regions using rapid sequences of 100ms pure tones and used tonotopy to identify cortical fields (Bizley et al., 2005). We specifically looked for the frequency tuning inversion: high-low-high moving in an antero-lateral direction, which marks the boundary between primary (A1) and secondary (dPEG) fields. At tonotopically mapped sites, we performed acute recordings with 64-channel integrated UCLA probes (Du et al., 2011), digital head-stages (RHD 128- Channel, Intan technologies) and OpenEphys data acquisition boxes and software (Siegle et al., 2017).

Raw voltage traces were processed with Kilosort 2 (Stringer, Pachitariu, Steinmetz, Reddy, et al., 2019), clustering and assigning spikes to putative single neurons. The clusters were manually curated with Phy (Rossant et al., 2016). Units were only kept for analysis if they maintained isolation and a stable firing rate over the course of the experiment. Unit isolation was quantified as the percent overlap of the spike waveform distribution with neighboring units and baseline activity. Isolation > 95% was considered a single unit and kept for analysis. We further filtered neurons based on the reliability of their responses, requiring a Pearson’s correlation > 0.1 between PSTH responses to the same stimuli (10 repetitions, 20 Hz sampling) drawn from random halves of repeated trials.

### Evaluating significance of sensory context effects

To measure effects of sensory context on sound-evoked activity, spike times for each unit were binned at 20 Hz. Activity was normalized as a Z-score based on mean and standard deviation of single-trial spike rate across the entire duration of the recording (spontaneous activity and during sound presentation). We define a contextual instance as a probe sound preceded by a pair of context sounds. Experiments using N=4 distinct sound samples produced 50 distinct contextual instances, and experiments using N=10 sounds produced 550. For each contextual instance, we computed the difference in the response to the probe between the two contexts. To track context effects over time, we calculated this difference at every 50-ms time bin (Δ Z-score).

To evaluate the significance of differences in the time-varying response, traditional analysis might use a T or U test for the response difference at every time bin. However, this approach leads to the problem of multiple comparisons and reduced sensitivity if using the Bonferroni corrections across many time points in the probe response. To maximize statistical power, instead, we used a cluster mass quantification of significance (Maris & Oostenveld, 2007), which corrects for multiple comparisons in the time domain, without sacrificing sensitivity.

For each contextual instance, we first calculated the T-score during each time bin of the probe response between each context. We then found groups of one or more contiguous time bins with significant T-scores of the same sign (p<0.05). Each of these groups defined a cluster with an associated score computed as the sum of the T-score for all time bins in the cluster. Finally, each cluster was assigned a p-value calculated by comparing the cluster score to its null distribution. This null distribution was obtained by calculating the maximum cluster statistic value (following the same procedure as above) for 11000 random shuffles of the context identity (Figure 1 F, J). We calculated this cluster-mass T-score and p-value for all contextual instances for a given neuron. We used the Bonferroni method to correct for multiple comparisons across contextual instances with a family error of alpha=0.05.

### Amplitude and duration of context effects

The temporal profiles of the context dependent differences in firing rate were diverse and irregular; therefore, we avoided describing them with monotonic distributions, e.g., exponential decay. Instead, we quantified the amplitude of contextual differences as the integral of the absolute difference (∫ |Δ Z-score|) across significant time bins and their duration as the time of the last significant time bin (Figure 1 G, K).

### Cortical field, context similarity and identity effects

We performed categorical multivariate linear regression to quantify the dependence of the amplitude and duration of context effects on cortical region (A1, dPEG), context similarity (*Silence*, *Different*, *Same*) and identity (ferret *Vocalization*, *non-Vocalization*). The amplitude and duration metrics for each contextual instance were normalized by dividing by the grand mean amplitude or duration across all neurons and contextual instances, thus scaling them to the percent change relative to the base category. For the cortical field effect, a univariate linear regression was fit using A1 as the base category. For the context similarity effect, since each instance was comprised by two contexts that could belong to different categories, data was repeated as needed with multiple labels, e.g., the same context effect value for an instance with *Same* and *Different* contexts appeared twice, labeled as *Same* or *Different*. Similarly for vocalization effects, instances that had at least one vocalization as context were classified as *Vocalization*, therefore (rare) instances with two vocalizations were classified the same as those with one. A multivariate linear regression was then fit using the context similarity and identity variables and their interaction. Most contextual instances were comprised of two *Different* contexts, which were *non-Vocalizations*, so this combination was assigned as the base variable. Significance of the regressed coefficients was quantified with a T-test over the residuals of the regression. Linear regression, using Ordinary Least Square minimization, and significance statistics were calculated using the python package Statsmodels (Seabold & Perktold, 2010).

Because the distributions of context effect amplitude and duration were strongly skewed and non-Gaussian, we validated the significance of context type and region differences with a non-parametric ANOVA (Kruskal-Wallis) and pairwise Dunn post hoc tests. We excluded instances with both *Silence* and *Vocalization* contexts from this analysis to make clear the effects of context type and vocalization in isolation, due to the strong interaction between them.

### Sparse population coding analysis

For every neuron, the significant context effects at every contextual instance yielded a coverage of the context instances, i.e., all combinations of probes and context-pairs. To describe the total contextual coverage of a multi-channel recording we calculated the union of context effects across all the neurons in the site. For every contextual instance, we included the value with the highest amplitude amongst all neurons in the union. In this case, we also considered each neuron as a source of multiple comparisons and corrected for it alongside the prior correction for number of contextual instances. The best neuron for each site was defined as the one with the greatest contextual coverage, i.e., having the greatest number of contextual instances with significant context effects.

To describe context effects of population activity in a low dimensional space, we used Principal Components Analysis (PCA). We computed the PCA transformation matrix on trial-averaged responses to n-sound sequences for all neurons in a recording site. We used this transformation matrix to project single-trial responses onto the first PC (i.e., the PC that explained the most variance in the average evoked response). This projection was then used to calculate context effects and coverage, following the same procedure as for single neurons.

### Dense coding model and decoding analysis

To measure the context and probe information present in the entire population activity, we trained a different linear support vector machine (SVM) to classify the single-trial population response vector at each 20 Hz time bin of the probe response. The SVM was trained to classify either the preceding context or the current probe identity. Prior to decoding, features (neuron activity) were normalized by removing the mean and scaling to unit variance. Performance was evaluated as the accuracy of label assignment, using 4-fold cross validation. SVMs were fit and evaluated using the scikit-learn python package (Pedregosa et al., 2011).

To simulate a dense code, the single unit data was transformed by equalizing context effects across all neurons, while preserving the mean probe response and trial-to-trial variability of each individual neuron. We define the response of neuron *i* to probe *p* following context *c* on trial *j* as *r*_*i,c,p,j*_(*t*). Each neuron’s average response to a probe was computed by averaging across context and trial,

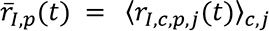

The average effects of contexts on every neuron-probe combination were computed by subtracting the average probe response and averaging across trials.

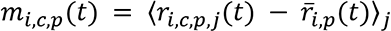

The population modulation by each context for each probe was computed as the L2 norm of the population activity vector 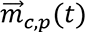 defined from *m*_*I,c,p*_(*t*). The differences in individual neuron activities were equalized by rotating each population vector towards the closest point in the identity diagonal, done by subtracting the difference of each neuron activity from the appropriate L2 fraction for *I* neurons,

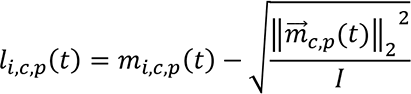

the vector rotations were applied to original single trial response, thus moving population context-trials around their corresponding probe average responses.

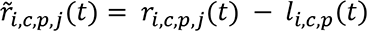

These vector rotations alter the response variance, particularly the fraction of variance coming from the context effects distribution around each probe, therefore we calculated this source of variance for the original and rotated responses,

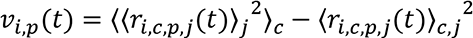

we then extracted from the rotated response the part corresponding to the context effects by averaging out probe and trial information,

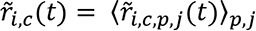

And corrected to the original variance by scaling the context effects, before adding them back.

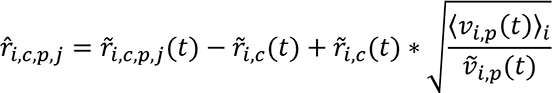

The decoding analysis was then applied to the transformed response *r̂*_*i,c,p,j*_(*t*), using the same approach as for the original data.

### Viral injection

For one animal, we injected auditory cortex with an adeno-associated virus serotype 2 (AAV2) containing channelrhodopsin 2 (ChR2), and mCherry under the inhibitory interneuron specific promote mDlx (Dimidschstein et al., 2016). Two craniotomies were drilled over adequate injection sites spanning A1. The responsiveness of injection sites was validated with electrophysiology. The injections were performed under Ketamine-Xylazine anesthesia and vitals were tracked through the procedure. The animal head was fixed using the previously implanted headcap. A glass injection needle with a beveled tip of ∼30 µm diameter, was coupled with flexible tubing to a 100µl syringe (Hamilton, 7656-01), controlled with an automated injector (New era pump systems, NE-1000). The syringe was preloaded with mineral oil (Sigma-Aldrich), which was used to prime the whole hydraulic system. 10µl of virus were front loaded, and ∼5µl were injected in each craniotomy. The injections were performed at a depth of ∼1.5mm, roughly in the middle of the cortical depth. To improve the coverage of the viral injection, we used a convection enhance delivery strategy (Weiss et al., 2020), where the delivery rate started from 0.5µl/min and was incremented by 0.5µl/min every 3 minutes until the desired volume was injected. The incubation period between injections and photo stimulation was 2 weeks.

### Immunohistochemistry

One animal was euthanized by pentobarbital overdose and transcardially perfused with 4% paraformaldehyde (PFA), the brain was extracted and placed in 4% PFA overnight. A ∼1.5 cm coronal section containing the injection sites cut. A high concentration protein jelly (BSA 30%, gelatin 0.5% w/w in DI-water) was mixed with 37% formaldehyde and 5% glutaraldehyde (200 µl and 80 µl respectively for 1 ml of protein jelly), and poured, quickly polymerizing into a gel block embedding the brain section. The embedded brain was cut into 100 µm coronal slices using a vibratome (Leica VT100S) and the slices containing the injection sites were screened by the faint mCherry residual red fluorescence.

The selected slices were stained with primary antibodies against GAD67 (mouse, MAB5406 Sigma-Aldrich) and mCherry (goat, AB0040-200 Sicgen). Secondary antibodies were Alexa 488 (Green) donkey anti mouse (715-545-150 Jackson Immunoresearch) and Cy3 (Red) donkey anti goat (705-165-147 Jackson Immunoresearch). Slices were permeabilized (permeabilization solution: PBS with 0.1% Triton x100, 1% BSA, 1% fish gelatin, 2% normal donkey serum) at room temperature (RT) for 2 hours, and then washed (PBS at RT for 10 minutes, 3 times). The slices were incubated with the primary antibodies at 4C for 48 hours, washed like before, incubated with the secondary antibodies for 48 hours at 4C and washed again. Antibodies were diluted in 1:1 permeabilizing solution and PBS, at dilutions of 1:500, 1:1000 and 1:2000 for primary, and 1:1000 for secondary. The staining was fixed by incubating with 4% PFA for 15 minutes at RT, followed by a final wash, and mounting in slices with Permount (Fisher Chemical). Images were acquired on an inverted Zeiss LSM 900 system confocal at the Advanced Light Microscopy core at OHSU, with the assistance of their staff. Image processing was done in FIJI, including gamma adjustment for the green channel (g=1.8).

### Optotagging

To photo stimulate and record from neurons simultaneously we attached an optic fiber (24 mm, 0.66 NA, 400 µm inner diameter, 430 µm cladding, 1.25 zirconia ferrule. Doric lenses MFC_400/430- 0.66_24mm_ZF1.25(G)_FLT) to UCLA 64 channel probe using nonconductive, encapsulating epoxy (Resinlab EP965). The optic fiber laid parallel and in contact with the probe shank, leaving ∼1.5mm of clearance between the fiber face and the electrodes. This clearance was enough so the array could be introduced into the cortex, and the optic fiber would lay on top of, or close to the dura. the ferrule was connected to a laser (Ikecool, IKE-473-100-OP) with an optic patch cord (2.5m, 0.57na, 400 µm inner diameter, 430 µm cladding. Doric lenses MFP_400/430/3000-0.57_2.5mm_FC-ZF1.25). Laser power delivered close to the dura was calibrated between 200-250mW/mm2.

The photo-stimulation consisted of 40 trials of a single 20ms flash delivered during silence. The inter trial interval was 1s, and the flash trials were randomly interspersed with control trials with no light. The light stimulation generated a significant photoelectric artifact consisting of high amplitude and low duration on and offset transients, and a sustained lower amplitude noise. The transients were eliminated by removing the 2ms right after laser on and off and interpolating to fill missing values. The lower amplitude ongoing noise was reduced by subtracting the common average of laser trials. Preprocessed data was then spike sorted as before, and the remaining artifacts appearing as spike clusters or outlier false-positive spikes on good cluster were discarded. Neurons were considered optotagged if they responded within 5ms to the light onset, with a train of action potentials reliable across trials (Figure 5a).

### Spike wave form analysis

Neurons were classified based on their average spike waveform width, which was calculated as the time between the depolarization valley and the hyperpolarization peak (Trainito et al., 2019). The spike width was calculated for all neurons with amenable waveforms (inverted mostly positive waveforms with multiple inflections, associated with axonal spikes (Barry, 2015; Sibille et al., 2022) were difficult to interpret and excluded). The distribution of spike widths followed a clear bimodal distribution. The width threshold was defined as the minimum of the kernel density estimation valley, and a safety range of 0.1ms around the threshold was kept as unclassified.

### Classified cell types comparisons

Comparisons of context amplitude, duration and coverage between narrow, broad spiking and optotagged neurons, were done with independent Kruskal Wallis tests for each metric, followed with Dunn post hoc tests with Bonferroni corrections. Amplitude and coverage were reported and plotted with medians, duration was reported with mean for clarity due to its discrete nature, poorly displayed by the median. The selection of summary statistic had no influence over the results of the statistical tests, and served only a display clarity purpose

### Pupillometry

To obtain a measure of global changes in arousal, pupil was recorded during experiments with a video camera (Adafruit TTL Serial Camera 397, M12 Lenses PT-2514BMP 25.0 mm) placed 10 cm from the eye. An infrared light was used to improve the contrast of the image. The pupil was kept partially contracted with an ambient light set to ∼1500 lux at the eye being recorded, this increased the dynamic range of pupil size. The pupil size was measured offline using software detailed in (Schwartz et al., 2020). Trials were classified by the median pupil size yielding a balanced number of large and small trials. This classification was performed independently for each contextual instance, calculating the median pupil size across the time interval containing both context and probe.

The context-independent pupil-dependent firing rate (mean Z-score) was calculated for all combinations of neurons and probes, averaging across contexts (Supplementary Figure 2A), and over the 1s probe duration. Nonresponsive neuron-probes (mean Z-Score<0.1) were filtered out for further analysis. The pupil-dependent context effects (mean Δ Z-score) was calculated for significant contextual instances identified previously using the cluster mass analysis. The magnitude of the pupil dependent effect was computed by averaging the Δ Z-scored context effects across the entire probe response. This metric was also calculated for non-overlapping 250ms time intervals (i:0-250, ii:250-500, iii:500-750, iv:750-1000). Values with low firing rate (mean Z-score<0.1) and small context effects (mean Δ Z-score<0.3) were filtered out. Prior to computing pupil effects, the (arbitrary) sign of context effects was flipped to be positive to enable pooling across all data.

A pupil modulation index (MI) was calculated for firing rates and context effects. MI was defined as (*large* − *small*) / (|*large*| + |*small*|), where |·| denotes the absolute value. Absolute value prevented numerical instability when the denominator approached zero. MI close to zero indicates no pupil effects, while negative and positive values indicate an increase for small and large pupil respectively.

### Model architecture

We trained encoding models to predict the activity of a neuron as a function of sound stimuli, its own past activity and that of other neurons in the population. All models followed the same Generalized linear model architecture:

Input sound was transformed into a log-spaced, 18-channel spectrogram (approximately 1/3 octave per channel) with amplitude log compression emulating cochlear dynamics (Pennington & David, 2022). Stimulus and spike signals were binned at 100 Hz. Spike data was averaged across trials and normalized to the peak value for each neuron. We used a standard linear-nonlinear spectro-temporal receptive field (STRF) as a baseline model of sensory encoding. The STRF spanned the 18 spectral channels over a window 300ms (30 10-ms bins) before the neuron response. A predicted response was computed by treating the STRF as a filter, convolving with the stimulus spectrogram in time and summing across spectral channels (Figure **6** A, top STRF).

To model the effect of the past neuronal activity, the response of all recorded neurons (including the neuron being predicted) was read over a time window spanning150-300ms before the neural response (15 10-ms bins). Neuronal activity was averaged over this time window, and the weighted sum of these averages was added to the STRF output (Figure 6A, past neuron, and population activity). Using the neuronal activity time average instead of every time point reduced the number of parameters, making the model more interpretable, where every parameter is an average synaptic strength in the population.

Traditional STRFs of 150ms can best capture the auditory driven responses in A1 and dPEG (Atiani et al., 2014), which happen in that time regime. Therefore, we used the population filters that were offset 150ms into the past to avoid capturing correlated sound evoked activity in the other neurons, and rather focus on the effect of recent population activity, i.e., a proxy for network activity. To disambiguate changes in model performance due to a longer temporal window extended by the population filter, we also extended the STRF to 300ms into the past, so its first half overlapped with the population filters. Ideally these “far past” sounds should carry little information about the sound evoked response, therefore, these weights tend to zero, and the nonzero weight of the STRF remain at short time lags (< 150ms). Finally, the summed output of the STRF and the neural filters was then passed through a rectified linear unit (ReLU) to account for spike threshold (Thorson et al., 2015).

We defined 4 different models based on this same architecture, by temporally scrambling different parts of their input: 1. a base STRF, achieved by scrambling both the self and population response 2. a “Self” model, where only the other neurons response was scrambled 3. a “Population” model (pop), where the self-response was scrambled 4. a Full model, with no scrambling. This scrambling effectively removes the predictive value of the scrambled data, while keeping the total number of model parameters unchanged. Temporal scrambling was done by independently shifting neuronal responses by random tens of seconds. We chose to shift instead of shuffling the data to keep the short-term temporal structure of the data and prevent the models from fitting to the grand average of the scrambled predictors.

All models were implemented using NEMS (Pennington & David, 2022), a flexible and readily available software developed in the lab. Optimization was performed using the ADAM gradient descent algorithm (Kingma & Ba, 2017).

### Model performance quantification

Model performance was quantified as the prediction correlation, computed as Pearson’s R across a predicted response assembled from N cross-validated model estimates (N=4 or 10, the number of different stimulus sequences).

To further quantify the ability of the different models to account for long-lasting context effects, we compared the context driven difference for neuronal responses and model predictions in each contextual instance. For the predictions, context effects were computed similarly to the ∫ |Δ Z-score| calculated from the actual response. However, there are two main differences. First, we used 0 to 1 normalization, instead of Z-scores, as models performed better this way. Second, there was no associated significance test (i.e., cluster mass analysis), due to the deterministic nature of our models.

We evaluated the accuracy with which models predicted context effects by calculating the Pearson correlation between the amplitude of context effects in the real data and those predicted by the model. This correlation was also calculated separately for 4 non-overlapping 250ms time intervals (i:0-250, ii:250-500, iii:500-750, iv:750-1000) spanning the probe response. This analysis was performed only on contextual instances that were significant according to the cluster mass test performed on the original data.

---

**Supplementary Figure 1.**
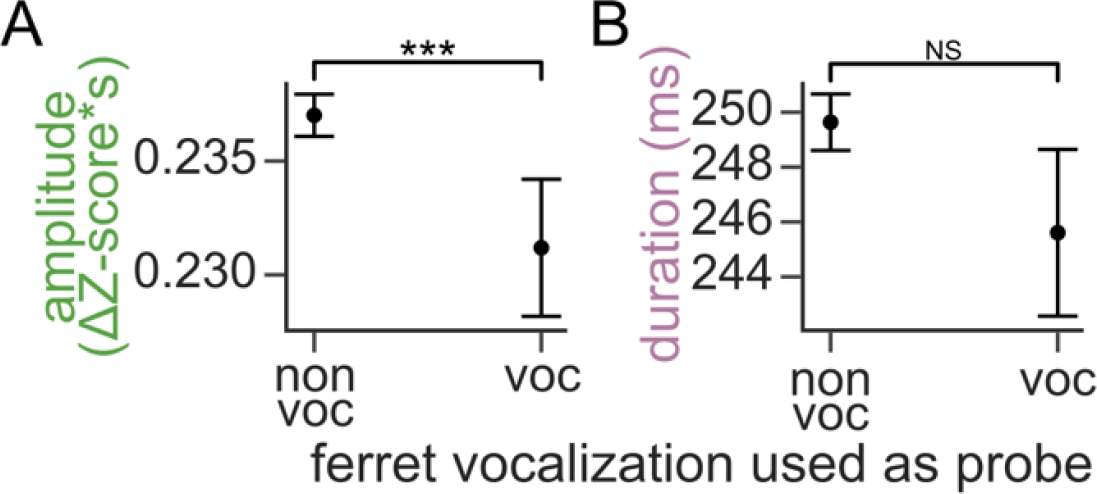
Weaker impact of context on responses to species-specific vocalization probe stimuli. A. Amplitude of context effects on a probe that was either a ferret vocalization or not (mean ± SEM: *non-vocalization*: 0.24 ± 0.0009, *vocalization*: 0.23 ± 0.0030 Z-score*s. Kruskal Wallis p<0.001). B. Same as in A for context effect duration (mean ± SEM: *non-vocalization*: 249.64±1.0281, *vocalization*: 245.61±3.0393 ms. Kruskal Wallis p=0.925).

**Supplementary Figure 2.**
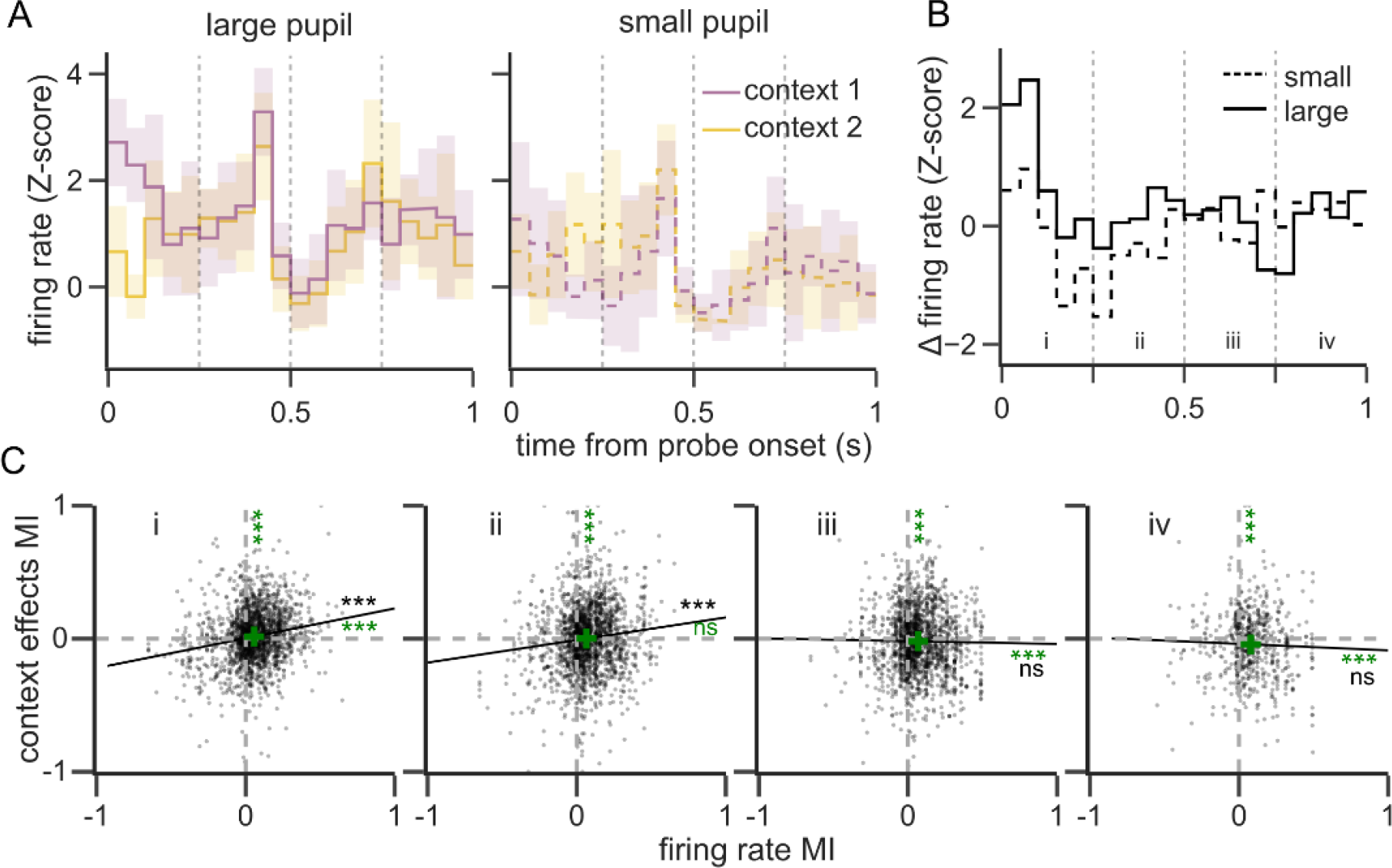
Large pupil is weakly correlated with decreased context effects. A. Example PSTH response of one neuron (line: mean, shade: SEM) to a probe sound following two contexts (color) for trials with large (left, solid lines) and small (right. dashed lines) pupil. When pupil is small, an overall reduction in firing rate reduces the amplitude of early context effects. B. Context-dependent difference (difference between context 1 and 2) for large (solid line) and small (dashed lines) pupil trials of the example in A. C. Comparison between pupil modulation index (MI: -1 to +1 for greater effects with small or large pupil, respectively) of firing rate (MI-fr) and context effect amplitude (MI-ce) for individual instances during 250ms intervals labeled in panel B (i to iv, n = 12868, 4942, 1815, 668, dots decimated for clarity). Vertical and horizontal dashed gray lines indicate MI=0 (no pupil effect). Green crosses indicate means on x and y. Significance of mean difference from zero on x and y is indicated with green symbols on the top and right sides (one sample T-test). Black lines indicate linear regression, with associated significance indicated in black (Wald test for nonzero slope). On average, firing rate was increased with large pupil during all intervals (mean MI-fr: i: 0.057; ii: 0.065; iii: 0.069; iv: 0.079. p<0.001 for all time intervals), while large pupil was associated with greater context effects during earlier intervals and smaller context effects during later intervals (mean MI-ce, p-value: i: 0.016, <0.0; ii: 0.001, 0.7; iii: -0.019, <0.001; iv: -0.042, <0.001). Dependence of firing rate and context effects on pupil size were significantly correlated during early intervals but not late intervals (Pearson’s R, p-value: i: 0.17, <0.001; ii: 0.11, <0.001; iii: -0.01, 0.5; iv: -0.03, 0.3). Statistical significance symbols, ***: p<0.01, ns: non-significant.

**Supplementary Figure 3.**
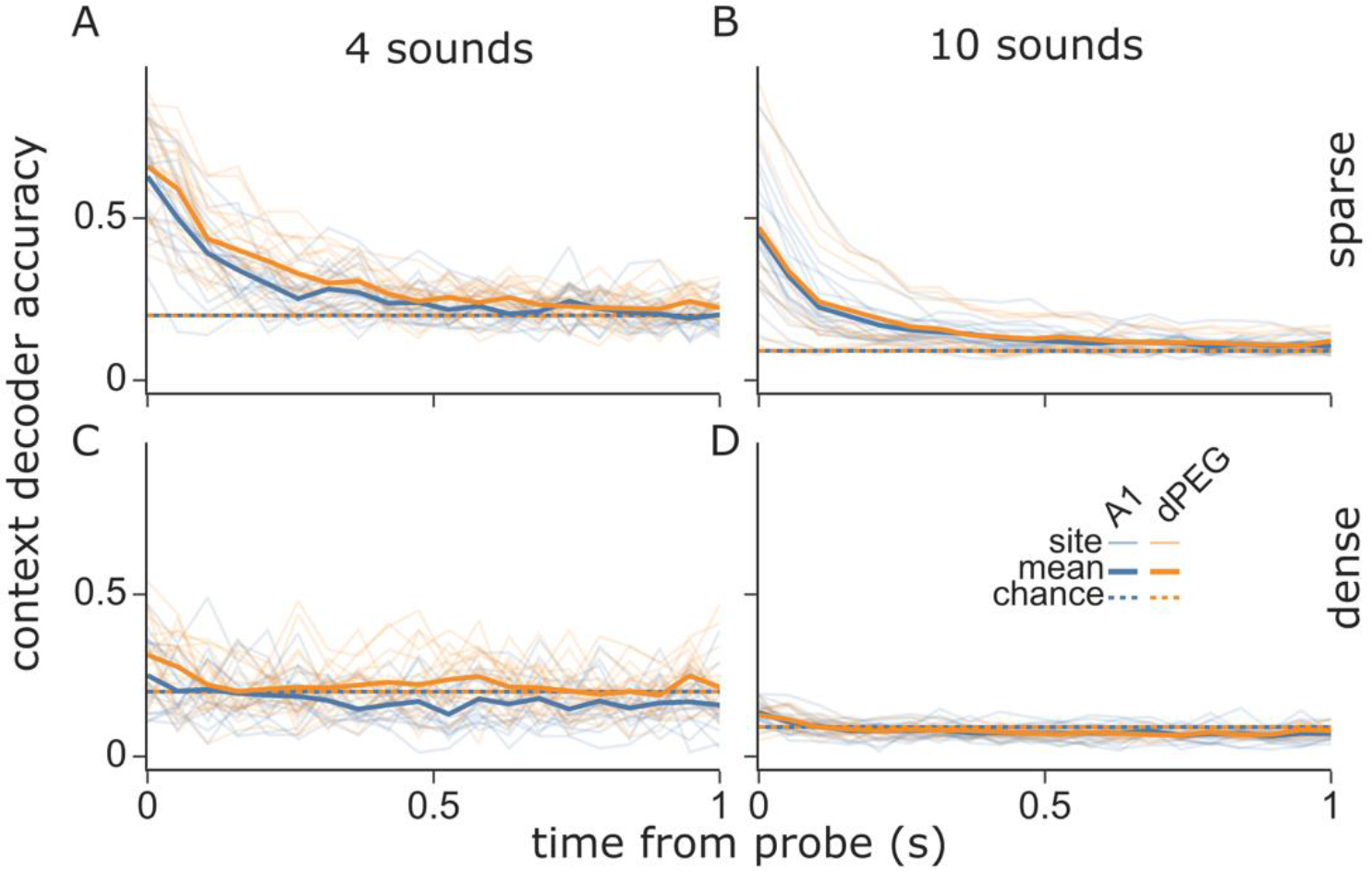
Decoding accuracy for different cortex regions and sound sets. A. Support vector machine accuracy at decoding the context identity from the population activity of sites (thin lines) and their mean (thick lines), recorded at A1 (blue) and dPEG (orange), subject to stimuli composed of 4 different sounds (5 contexts, including silence). Chance performance is indicated with dotted lines. B. Same as in A but for stimuli composed of 10 different sounds (11 contexts). C-D. Same as in A-B but after transforming the data to impose a dense representation of the context information.

**Supplementary Figure 4.**
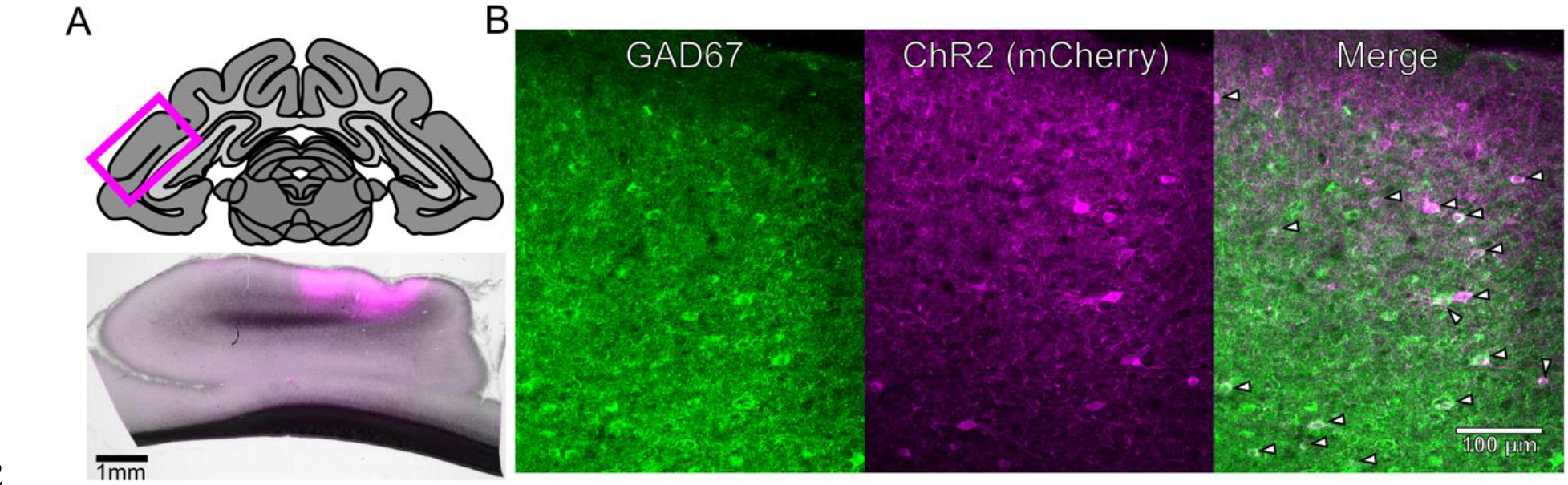
Validations of Channelrhodopsin-2 expression in inhibitory interneurons. A. Top: schematic of a coronal section of the ferret brain encompassing a section of the ectosylvian gyrus and A1, indicated by the magenta box. Bottom: bright field image with superimposed red fluorescence of the virally transduced cell (mCherry, magenta) around the injection sites on the ectosylvian gyrus indicated above. Scale bar: 1mm. B. immunohistochemistry of the injection site in A, showing GAD67 positive inhibitory interneurons (green), transduced cells expressing ChR2 alongside the reporter mCherry, under the inhibitory interneuron specific promoter mDlx (magenta), and the channel merge with arrows indicating neurons co-expressing the markers. Most transduced neurons are inhibitory interneurons. Scale bar: 100 µm.

